# Granzyme A inhibition reduces inflammation and increases survival during abdominal sepsis

**DOI:** 10.1101/2020.06.05.135814

**Authors:** Marcela Garzón-Tituaña, José L Sierra-Monzón, Laura Comas, Llipsy Santiago, Tatiana Khaliulina-Ushakova, Iratxe Uranga-Murillo, Ariel Ramirez-Labrada, Elena Tapia, Elena Morte-Romea, Sonia Algarate, Ludovic Couty, Eric Camerer, Phillip I Bird, Cristina Seral, Pilar Luque, José R Paño-Pardo, Eva M Galvez, Julián Pardo, Maykel Arias

## Abstract

Sepsis is a serious syndrome characterised by a dysregulated systemic inflammatory response. Here we have analysed the role and the therapeutic potential of Granzyme A (GzmA) in the pathogenesis of peritoneal sepsis using the Cecal Ligation and Puncture (CLP) polymicrobial sepsis model and samples from humans undergoing abdominal sepsis.

Elevated GzmA was observed in serum from patients with abdominal sepsis suggesting that GzmA plays an important role in this pathology. In the CLP model GzmA deficient mice, or WT mice treated with an extracellular GzmA inhibitor, showed increased survival, which correlated with a reduction in proinflammatory markers in both serum and peritoneal lavage fluid. GzmA deficiency did not influence bacterial load in blood and spleen indicating that GzmA has no role in bacterial control. Mechanistically, we found that extracellular active GzmA acts as a proinflammatory mediator in macrophages by inducing the TLR4-dependent expression of IL-6 and TNFα.

Our findings implicate GzmA as a key regulator of the inflammatory response during abdominal sepsis, and suggest that it could be targeted for treatment of this severe pathology.

## INTRODUCTION

Peritonitis is an inflammation of the peritoneum, usually following infection of the serous membrane that covers the abdominal cavity and the organs contained therein (Brown & Caballero Alvarado, 2019). Peritonitis is classified as primary, secondary or tertiary depending on disease severity and effect on the different organs (Muresan *et al*, 2018). Primary peritonitis is associated with undamaged intra-abdominal cavity organs. The most frequent forms are spontaneous bacterial peritonitis, associated with advanced liver disease (infected ascites), and infection in patients undergoing peritoneal dialysis. Secondary peritonitis is an infection in the peritoneal cavity due to a perforation of the gastrointestinal tract by ulceration, ischemia or obstruction; a post-surgical infection or a closed or penetrating trauma. Tertiary peritonitis is a persistent intra-abdominal infection after at least 48 hours of adequate therapy of primary or secondary peritonitis (Muresan *et al.*, 2018). Peritonitis is a leading cause of sepsis (Brown & Caballero Alvarado, 2019), a serious syndrome characterized by a dysregulated systemic inflammatory response (Gotts & Matthay, 2016; Iskander *et al*, 2013).

Granzymes (Gzms) are a family of serine proteases that, together with perforin, constitute the main components of the cytotoxic lymphocyte granule exocytosis pathway. Gzms are classified according to their cleavage specificity. Five Gzms in humans (A, B, H, K, and M) and ten in mice (A, B, C, D, E, F, G, K, M and N) have been described (Arias *et al*, 2017). The granule exocytosis pathway is a specialized form of intracellular protein delivery by which lymphocytes release perforin and Gzms into infected or compromised cells. Perforin creates short-lived pores in the plasma membrane of target cells allowing the passage of Gzms into the cytosol of target cells where they carry out their effector functions (Wensink *et al*, 2015). Gzms can also be released into the extracellular milieu where they regulate different extracellular processes independently of their ability to induce cell death (Arias *et al.*, 2017; Turner *et al*, 2019), Traditionally, it was assumed that all Gzms acted as cytotoxic proteases. However, recent evidence suggests that only GzmB has clear cytotoxic capacity, while the cytotoxicity of others such as GzmA and GzmK is controversial (Cullen *et al*, 2010; Chowdhury & Lieberman, 2008; Joeckel & Bird, 2014; Martinez-Lostao *et al*, 2015; Voskoboinik *et al*, 2015; Wensink *et al.*, 2015).

It has been recognised that some Gzms like GzmA may act as pro-inflammatory mediators contributing to the pathophysiology of different inflammatory disorders including rheumatoid and viral arthritis (Santiago *et al*, 2016; Wilson *et al*, 2017), colitis (Tew *et al*, 2016), endotoxicosis (Anthony *et al*, 2010; Metkar *et al*, 2008) or bacterial sepsis (Arias *et al*, 2014; van den Boogaard *et al*, 2016). Furthermore, *in vitro* studies have reported that GzmA induces the expression of pro-inflammatory cytokines in several cell types including monocytes, macrophages, endothelial and epithelial cells and fibroblasts (Campbell *et al*, 2018; Joeckel *et al*, 2011; Metkar *et al.*, 2008; Sharma *et al*, 2016; Sower *et al*, 1996). However, the role of GzmA in abdominal sepsis, the second most common form of sepsis in humans, is not known.

This study aimed to analyse the involvement of GzmA in abdominal sepsis, and gain insight into its therapeutic potential. We examined serum samples from patients with intraabdominal sepsis; and investigated the role of GzmA in sepsis using the Cecum Ligation and Puncture (CLP) mouse model, which best mimics the septic response during human peritonitis (Lewis *et al*, 2016).

## RESULTS

### Extracellular GzmA is increased in patients undergoing abdominal sepsis

We analysed the levels of GzmA in serum from patients with abdominal sepsis, who were diagnosed with peritonitis with a SOFA score >2, and compared with the levels in serum from healthy donors. Patients with peritoneal sepsis presented elevated levels of GzmA at admission and for 72h thereafter that were significantly above healthy donor levels both at admission and after 24h (Figure 1). This shows that serum GzmA is elevated during abdominal sepsis and suggests that extracellular GzmA could be involved in its pathogenesis.

**Figure 1.**
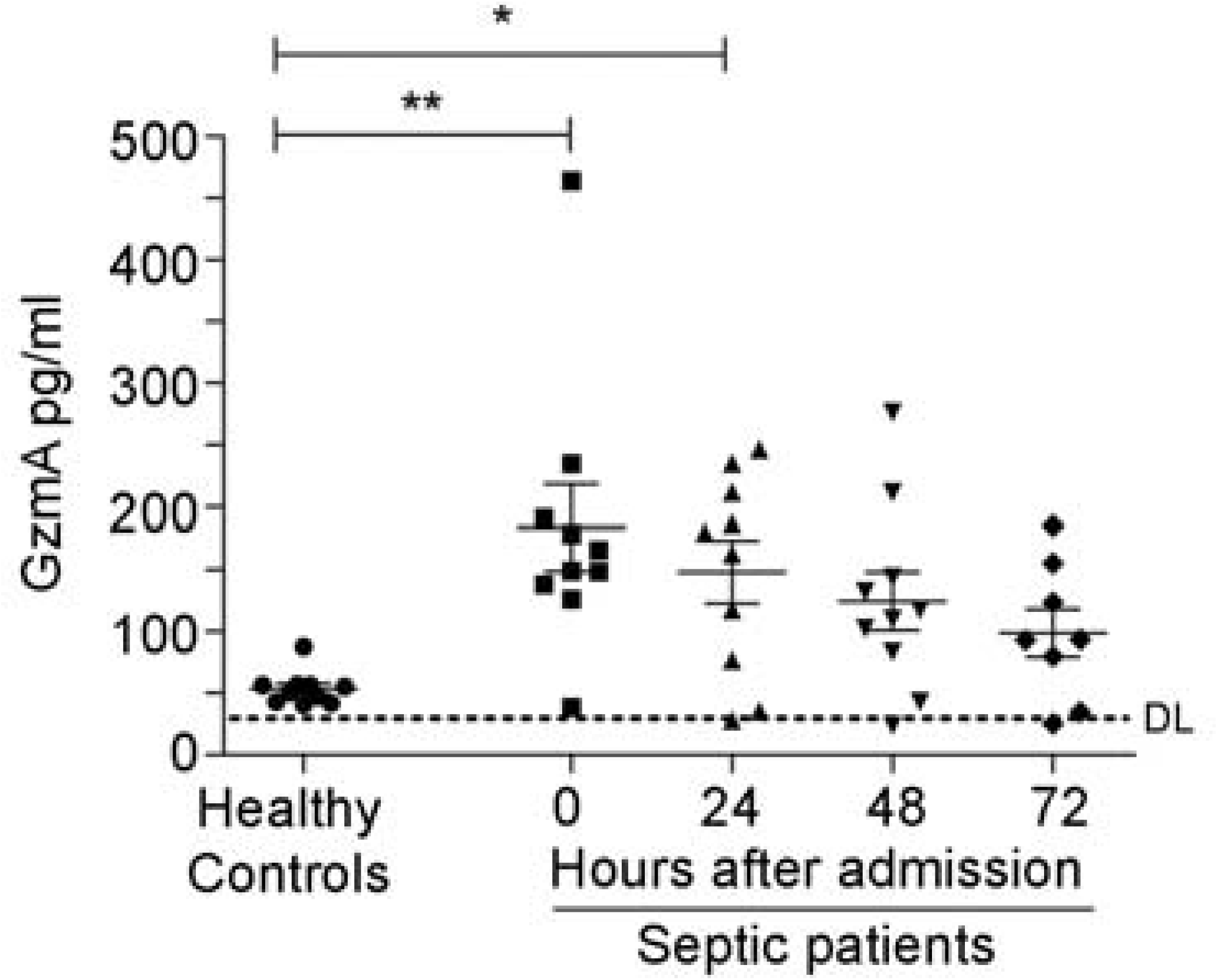
Increased levels of extracellular GzmA are observed in patients with abdominal sepsis. Serum levels of GzmA were analysed by ELISA in healthy donors (n=10) and compared with GzmA levels from patients with a diagnosis of abdominal sepsis (n=10) at admission and during the first 72 h. Statistical analyses were performed by one-way ANOVA test with Bonferroni’s post-test *P < 0.05; **P < 0.01.

### *GZMA* deficient mice are protected from CLP

Once we had observed increased levels of extracellular GzmA in patients with abdominal sepsis, we decided to analyse the relevance of these findings by testing the role of GzmA in the CLP mouse model, one of the murine models that better mimics the complex septic response in human abdominal sepsis (Lewis *et al.*, 2016). A severe sepsis CLP protocol was applied to WT and *GZMA*^-/-^ mice as indicated in methods. In order to mimic the clinical management of septic patients, a broad-spectrum antimicrobial treatment was administered starting six hours after CLP, and survival monitored for 14 days.

As shown in figure 2 survival of *GZMA*^-/-^ mice was significantly higher than WT controls. Less than 40% of WT mice survived CLP, while more than 70% of *GZMA*^-/-^ mice were still alive after 2 weeks. As expected, all sham mice survived, which confirms that the surgical technique was performed effectively. These results indicate that GzmA plays an important role during severe abdominal sepsis induced by CLP *in vivo*.

**Figure 2.**
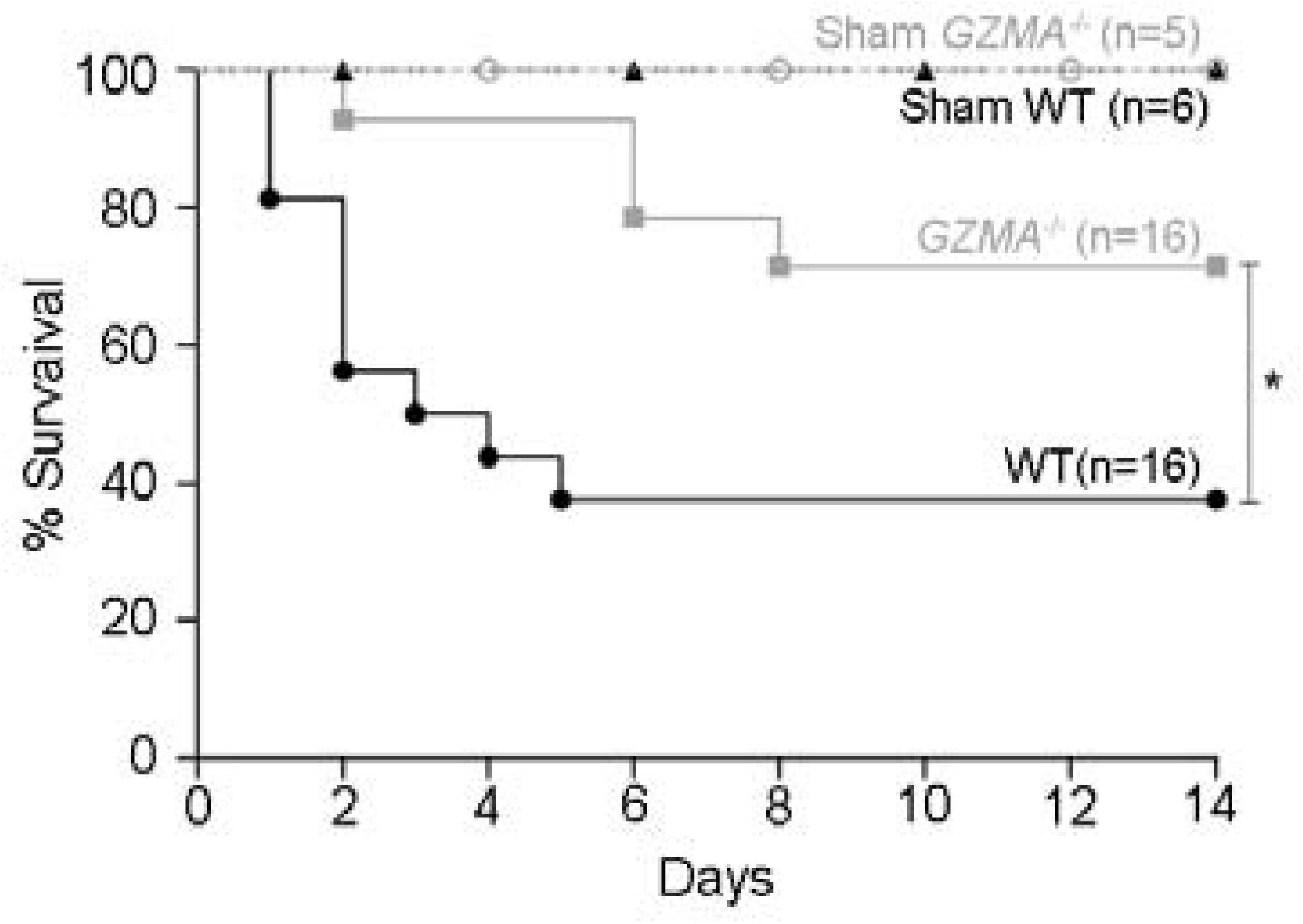
The absence of granzyme A increases survival in mice during sepsis induced by CLP. Sepsis was induced by CLP in WT and *GZMA*^-/-^ mice as described in materials and methods. After 6 h, a mixture of antibiotics, ceftriaxone (30 mg/kg) + metronidazole (12.5 mg/kg) was administered every 24 h for 5 days. WT and *GZMA*^-/-^ sham operated mice underwent the same procedure but without the ligation and puncture of the cecum. Survival was monitored for 14 days. The data correspond to the indicated number of biological replicates (individual mice) from three independent experiments. Statistical analysis was performed using logrank and Gehan-Wilcoxon test. *p < 0.05.

### *GZMA* deficient mice show reduced levels of proinflammatory cytokines during abdominal sepsis

Next, we analysed if the protection observed in *GZMA* deficient mice during sepsis induced by CLP correlated with reduced serum level of proinflammatory cytokines. Thus, we monitored the concentrations of IL-6, IL-1α, IL-1β and TNFα in plasma and peritoneal lavage fluid (Figure 3). *GZMA*^-/-^ mice showed significantly lower levels of all cytokines in both plasma and peritoneal lavage fluid after 6 and/or 24h of CLP induction. It should be noted that although all cytokines were reduced in the biological fluids of *GZMA*^-/-^ mice after CLP, only IL6 and IL1α were significantly lower than controls at all time points in both plasma and peritoneal fluid. After 24h all cytokines were significantly reduced in septic *GZMA*^-/-^ mice. This confirms that GzmA regulates the generation of proinflammatory cytokines *in vivo* during abdominal sepsis.

**Figure 3.**
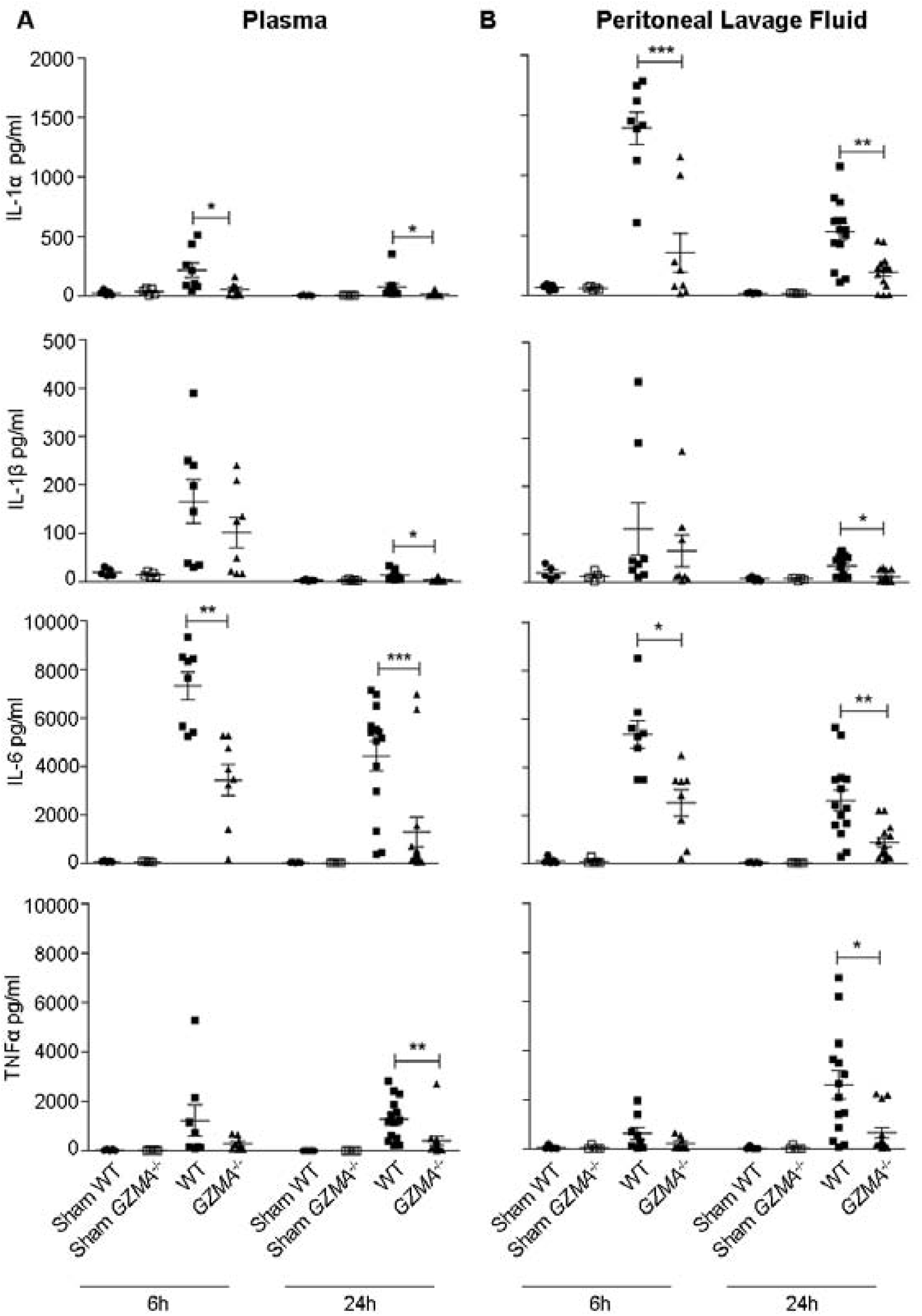
The absence of granzyme A reduces the level of proinflammatory cytokines during sepsis induced by CLP. Sepsis was induced by CLP in WT and *GZMA*^-/-^ mice as described in materials and methods. After 6 h, a mixture of antibiotics, ceftriaxone (30 mg/kg) + metronidazole (12.5 mg/kg) was administered every 12 h. WT and *GZMA*^-/-^ sham operated mice underwent the same procedure but without CLP. After 6 or 24 h of sepsis induction, mice were sacrificed and the levels of IL-1α, IL-1β, TNFα and IL-6 in plasma and peritoneal lavage fluids were determined by ELISA. Data are presented as mean ± SEM of 5 (sham) or at least 8 (CLP) different biological replicates (individual mice) from 3 independent experiments. Statistical analysis was performed by one-way ANOVA test with Bonferroni’s post-test *P < 0.05; **P < 0.01; ***P < 0.001.

### The absence of GzmA does not compromise anti-bacterial control during CLP

*GZMA* deficiency reduces the generation of inflammatory cytokines *in vivo.* Thus, it is possible that the reduced inflammatory response affects efficient clearance of local and/or systemic bacterial infection after CLP. We analysed the total aerobic bacterial load in blood, spleen, liver, lungs and peritoneal fluid during CLP-induced sepsis. As shown in Figure 4A, WT and *GZMA*^-/-^ mice exhibited similar bacterial loads in peritoneal lavage fluid, blood, liver, spleen and lung after 6 and 24 h of sepsis induction. Although total bacterial counts were similar in both mouse strains, it is possible that the control of specific bacterial species is compromised in the absence of GzmA. Thus, we characterised the bacterial species presented in the different organs and fluids and quantified their individual level by MALDI-TOF mass spectrometry. The analysis showed that the most frequent species present were *Lactobacillus murinus, Enterococcus faecalis, Staphylococcus sciuri, Streptococcus sp* and *Escherichia coli.* The bacterial load of these species in WT and *GZMA*^-/-^ mice was similar in peritoneal fluids, blood, spleen, liver and lung, after 6 and 24 h of sepsis induction (Fig. 4B). These results indicate that GzmA deficiency does not compromise control of aerobic bacteria during polymicrobial abdominal sepsis.

**Figure 4.**
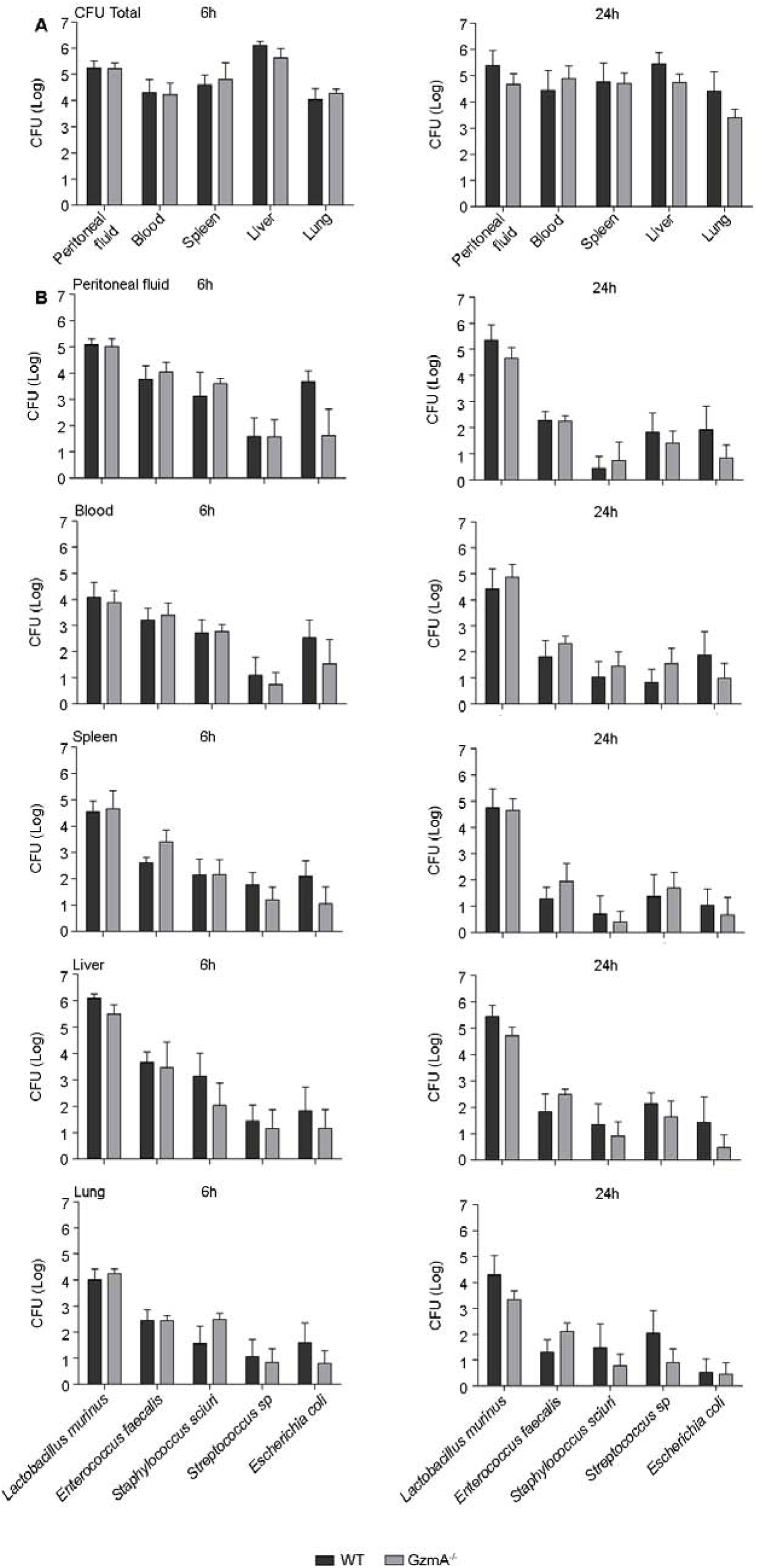
GzmA is not involved in the control of bacterial pathogens during sepsis induced by CLP. Sepsis was induced by CLP in WT and *GZMA*^-/-^ mice as described in materials and methods. After 6 h, a mixture of antibiotics, ceftriaxone (30 mg/kg) + metronidazole (12.5 mg/kg) was administered every 12 h. **A.** After 6 or 24 h of sepsis induction, mice were sacrificed and the total number of CFU from aerobic bacteria was determined in peritoneal fluids, blood, spleen liver and lung. **B.** The most abundant bacterial species were identified by MALDI-TOF mass spectrometry and the number of CFU of these species was determined in peritoneal lavage fluids, blood, spleen liver and lung. Data are presented as mean ± SEM from 5 biological replicates (individual mice) in each group.

### GzmA expression increases in NK cells during abdominal sepsis

Once we confirmed that the absence of GzmA increases the survival of mice during abdominal CLP-induced sepsis, correlating with a reduced inflammatory response that did not compromise bacterial control, we decided to establish the cellular source of GzmA. We focused on the major cell populations known to express GzmA in blood and spleen (Arias *et al.*, 2017): NK and T cells, including NKT cells. The gating strategy is summarised in Figure 5A. Cells were stained with NK1.1, CD3, CD4 and CD8 antibodies in order to distinguish NK cells (NK1.1^+^CD3^-^ cells) from the main T cell subsets, NKT (NK1.1^+^CD3^+^), CD4^+^ T (NK1.1^-^CD3^+^CD4^+^) and CD8^+^T (NK1.1^-^ CD3^+^CD8^+^) cells. As shown in Figure 5, GzmA expression was significantly enhanced in NK cells from septic mice in both spleen and peripheral blood. In contrast, the expression of GzmA in the main T cell subsets (NKT, CD8^+^ Tor CD4^+^T cells) was very low and did not increase during sepsis.

**Figure 5.**
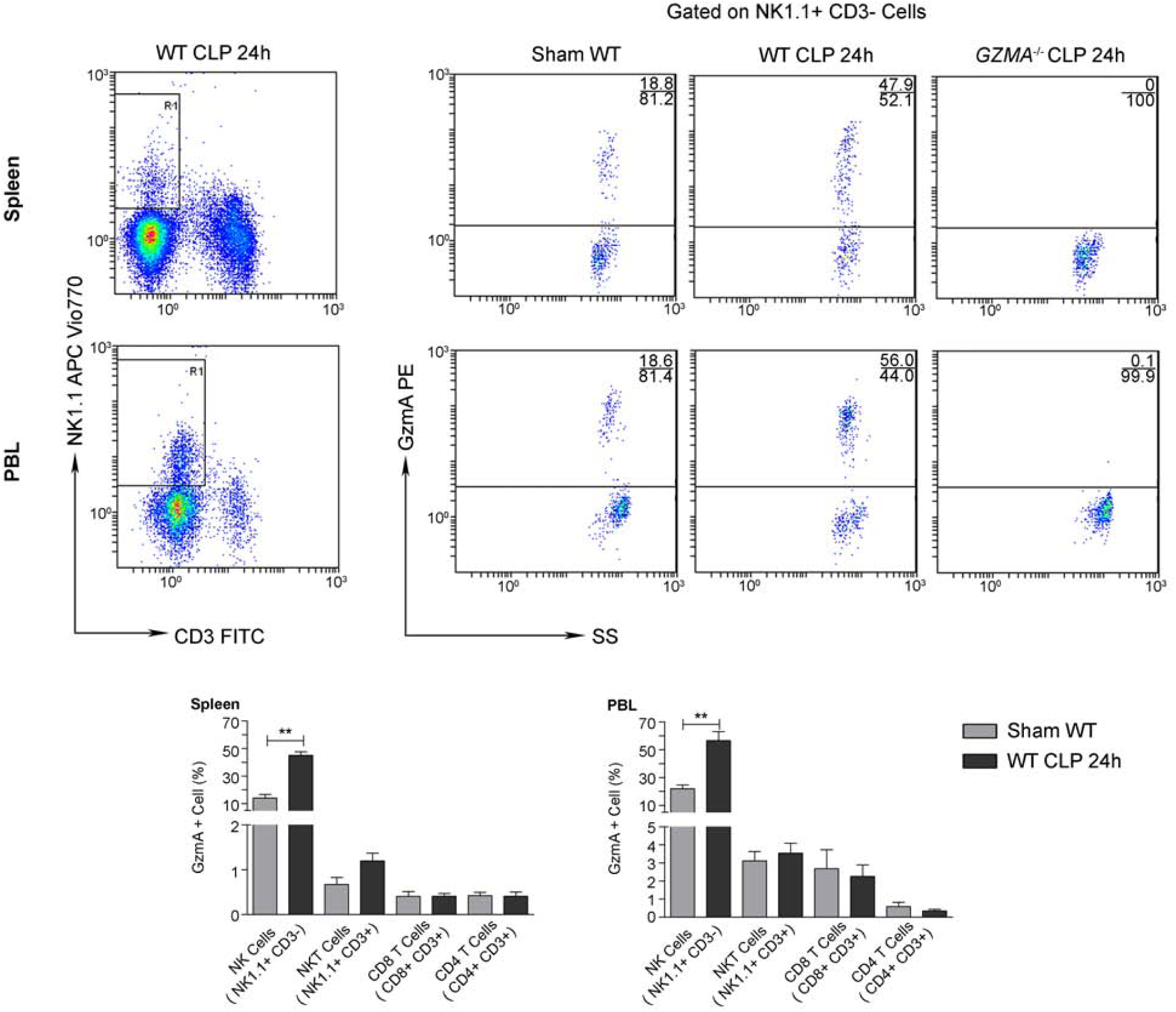
GzmA expression is increased in NK cells from septic mice. Sepsis was induced by CLP in WT and *GZMA*^-/-^mice as described in materials and methods. Sham WT operated mice underwent the same procedure but without the ligation and puncture of the cecum. After 24 h mice were sacrificed and the intracellular expression of GzmA on NK cells (NK1.1^+^CD3^-^), NKT cells (NK1.1+ CD3+), CD8+ T cells (CD8+ CD3+) and CD4+ T cells (CD4+, CD3+) was analysed in splenocytes and PBLs by flow cytometry. A representative experiment is shown via dot plot. Numbers show the cell percentage in each quadrant. Data in graphs represent the mean ± SEM of the percentage of GzmA positive cells in 4 biological replicates from 2 independent experiments. Statistical analysis was performed by unpaired student’s t test. **p < 0.01.

### Therapeutic inhibition of GzmA with Serpinb6b increases survival and reduces inflammation during CLP

Once the effect of GzmA absence was analysed using a genetically deficient model, we decided to further analyse the potential of using GzmA as therapeutic target to treat abdominal sepsis. To this aim we analysed the effect of a GzmA inhibitor on WT animals undergoing abdominal sepsis. We used serpinb6b, a recently described natural specific inhibitor of mouse GzmA, which does not inhibit other gzms (Kaiserman *et al*, 2014). After sepsis induction a group of WT and *GZMA* deficient mice were treated with the inhibitor and survival was monitored during 14 days. As shown in figure 6A, WT mice treated with antibiotics and serpinb6b showed a significant improvement in survival compared with WT mice treated with antibiotics only. Survival of septic WT mice treated with serpinb6b was similar to the survival of *GZMA* deficient mice. In contrast, the inhibitor did not have any effect on *GZMA* deficient mice suggesting that the effect observed in WT mice was specifically a consequence of GzmA inhibition. In parallel, a group of mice treated or not with the inhibitor were sacrificed and the level of IL-6 in plasma and peritoneal fluid was determined. In WT mice the therapeutic inhibition of GzmA significantly reduced the level of IL-6 in plasma and in peritoneal fluids. By contrast, the administration of Serpinb6b in septic *GZMA*^*-/-*^ mice did not reduce the levels of IL-6, again confirming the specificity of Serpinb6b to inhibit GzmA (Fig. 6B). These results confirm that GzmA inhibition in septic WT mice mimics the effect observed in *GZMA* deficient mice and provide a proof of principle that inhibition of GzmA could be useful for the treatment of abdominal sepsis.

**Figure 6.**
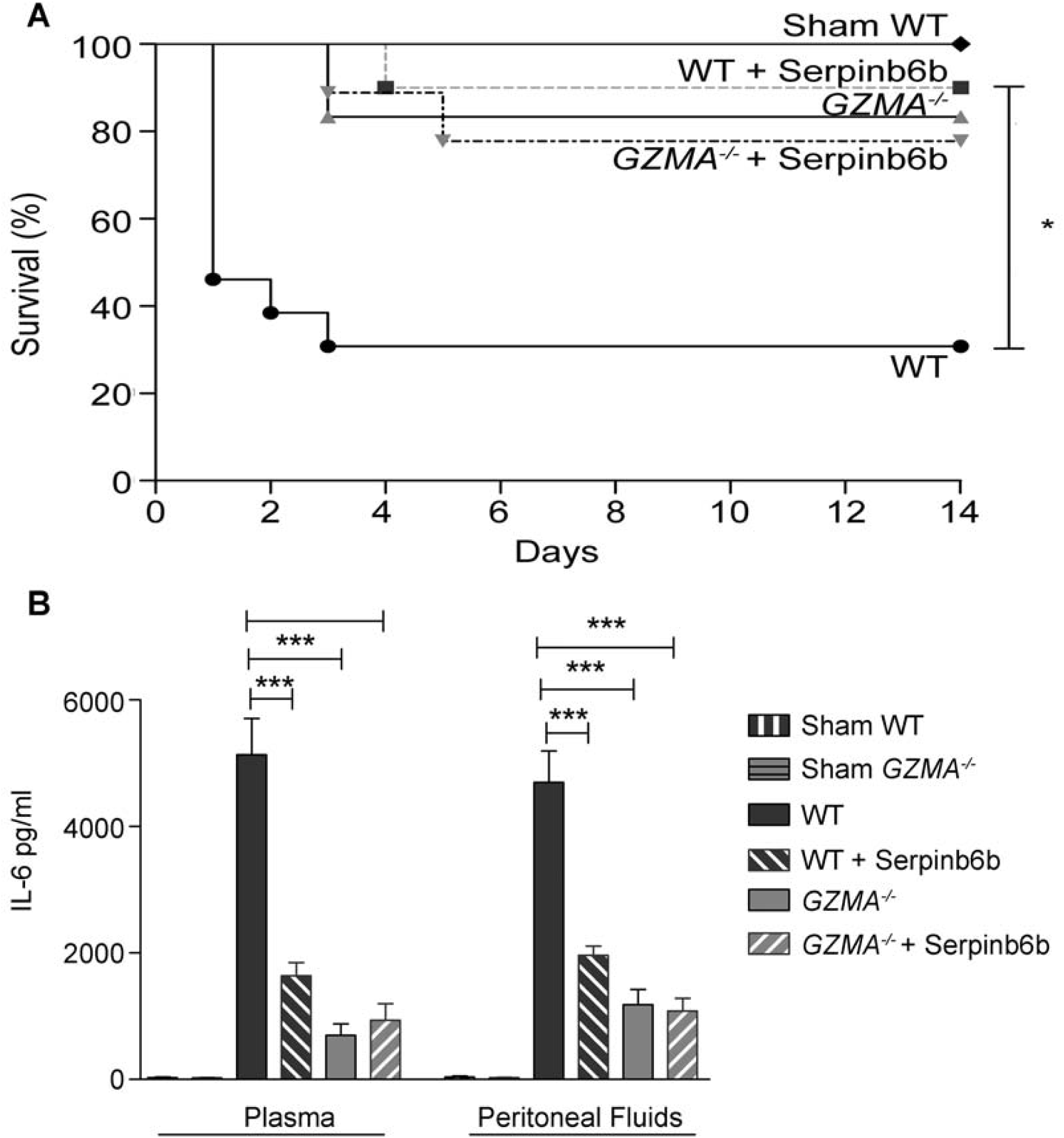
Inhibition of extracellular GzmA improves sepsis outcome and reduces inflammation. Sepsis was induced by CLP in B6 and *GZMA*^-/-^ mice as described in materials and methods. Immediately after surgery mice were treated with 40 µg of Serpinb6b in 100µl of PBS (10 WT and 9 *GZMA*^*-/-*^ mice). This treatment was repeated 6h later and once a day during 5 days. Control mice received only 100µl of PBS ip (13 WT and 6 *GZMA*^*-/-*^ mice). After 6 h a mixture of antibiotics, ceftriaxone (30 mg/kg) + metronidazole (12.5 mg/kg) was administered ip once a day for 5 days. WT sham operated mice (4 WT mice) underwent the same procedure but without the ligation and puncture of the cecum. **A.** Survival was monitored during 14 days. The data correspond to the indicated number of mice combined from two independent experiments. Statistical analyse was performed using logrank and Gehan-Wilcoxon test. *p < 0.05. **B.** 24 h after sepsis induction mice were sacrificed and the levels of IL-6 in plasma and peritoneal fluids was determined by ELISA. Data are presented as mean ± SEM of 4 (Sham) or 6 (CLP) biological replicates from 2 independent experiments. Statistical analysis was performed by one-way ANOVA test with Bonferroni’s post-test *P < 0.05; **P < 0.01; ***P < 0.001.

### The inflammatory response induced by GzmA on mouse macrophages depends on TLR4 expression

Increased levels of GzmA were found in patients with abdominal sepsis (Fig. 1) and human GzmA is known to induce the generation of inflammatory cytokines in monocytes and macrophages (Metkar *et al.*, 2008; Wensink *et al*, 2016). Thus, in order to find out how extracellular GzmA contributes to abdominal sepsis, we analysed the generation of inflammatory cytokines in mouse macrophages and the potential involvement of TLR4 in this process.TLR4 is a key receptor activated by microbial PAMPs like LPS and different endogenous ligands or DAMPs (Yu *et al*, 2010) and it has been previously found to be involved in abdominal sepsis (Daubeuf *et al*, 2007).

Thus, the effect of extracellular GzmA on the generation of IL6 and TNFα by M1 bone marrow-derived macrophages from WT and TLR4 deficient mice was analysed. We found that recombinant mGzmA produced in *E. coli* induced the expression of IL-6 on M1 WT macrophages which was reduced in absence of TLR4 (Fig. 7A). This result indicates that GzmA induces the expression of IL-6 by a mechanism dependent on TLR4.

**Figure 7.**
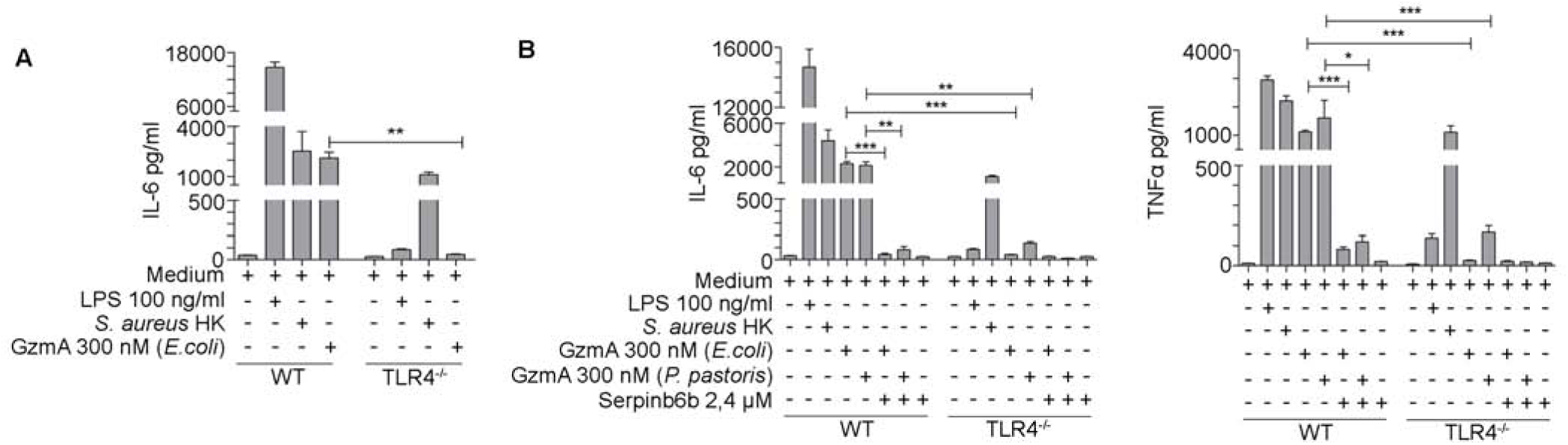
Active extracellular GzmA induces the expression of IL-6 and TNFα in macrophages by a mechanism dependent of TLR4. **A.** Macrophages differentiated from WT or TLR4^-/-^ mouse bone marrow cells were treated with active GzmA (300 nM) produced in *E.coli*, LPS 100 ng/ml or *S. aureus* heat killed (*S. aureus* HK,1 x10^6^ CFU/ml). After 24 h the supernatant was collected to determine the levels of IL-6 by ELISA. **B.** Macrophages differentiated fromWT or TLR 4^-/-^mouse bone marrow cells were treated with active GzmA (300 nM) produced in *E. coli* or in *P. pastoris*, GzmA inactivated with Serpinb6b, LPS 100 ng/ml or *S. aureus* HK (1 x10^6^ CFU/ml). After 24 h the supernatant was collected to determine the levels of IL-6 and TNFα by ELISA. Data are represented as the mean ± SEM of two independent experiments performed by duplicate. Statistical analyses were performed by one-way ANOVA test with Bonferroni’s post- test, *P < 0.05, **P < 0.01; ***P < 0.001.

It has been reported that inactive GzmA potentiates the proinflammatory effects of LPS in monocytes by a TLR4-dependent mechanism (Wensink *et al.*, 2016). In order to discount the potential contribution of LPS during inflammation induced by recombinant GzmA and to analyse if GzmA-induced inflammation is dependent on the enzymatic activity of this protease, we compared the inflammatory activity of active recombinant mGzmA produced either in *E. coli* or in *P. pastoris* on WT and TLR4^-/-^ macrophages; in addition, we used GzmA inactivated with Serpinb6b and the expression of IL-6, IL-1β and TNFα was quantified by ELISA (Fig. 7B). We observed that active GzmA produced from both sources induced similar levels of expression of IL-6 and TNFα in M1 WT macrophages. In addition, we observed that the expression of these proinflammatory cytokines was abolished when GzmA was inactivated with Serpinb6b or in absence of TLR4, indicating that the expression of IL-6 and TNFα induced by GzmA requires enzymatically active GzmA and is dependent on TLR4 (Fig. 7B). It is important to note that the potential confounding contribution of contaminating LPS in these experiments can be discounted because recombinant GzmA produced in *P. pastoris* was used.

## DISCUSSION

In this study we have analysed the contribution of GzmA to abdominal sepsis by investigating the presence of extracellular GzmA in serum from patients with peritoneal sepsis as well as the role of GzmA in the mouse model of CLP. GzmA was found to be elevated in human serum from septic patients. The relevance of these findings was supported by the observation that GzmA deficient mice were resistant to CLP. GzmA deficiency reduced inflammation without altering bacterial dissemination in the CLP model. A GzmA inhibitor enhanced survival and reduced inflammation in septic WT mice to a similar degree as observed with genetic deficiency, suggesting therapeutic potential in targeting GzmA in peritoneal sepsis.

One of the advantages of CLP model is that the pathogens are endogenous, mimicking the traumatic injury that leads to peritonitis in humans. In addition, CLP sepsis shows a high degree of similarity with the progression of human sepsis, displaying both the hyper- and hypo-inflammatory responses characteristic of human sepsis (Buras *et al*, 2005; Nemzek *et al*, 2008). It has been shown previously that GzmA might contribute to septic shock induced by LPS (Anthony *et al.*, 2010; Metkar *et al.*, 2008) or to sepsis induced by single bacterial agents like the mouse pathogen *B. microti (Arias et al., 2014), S. pneumoniae* (van den Boogaard *et al.*, 2016) or *E. coli* (Garcia-Laorden *et al*, 2017). However, the role of GzmA in abdominal polymicrobial sepsis had so far not been explored. Most importantly, the effect of GzmA inhibition in septic mice had not been analysed in any of these models. Our study thus adds to previous literature and confirms that GzmA is a key mediator of sepsis associated with different bacterial pathogens.

Our data also suggest that inflammation induced by GzmA plays a critical role in the development of sepsis during peritonitis. We observed lower production of IL-1α, IL-1β, IL-6 and TNFα in serum and in peritoneal fluids in GzmA deficient mice. All these cytokines play an important role in sepsis (Chong & Sriskandan, 2011; Linkermann *et al*, 2012; Matsumoto *et al*, 2018; Vanden Berghe *et al*, 2014). In addition, our results employing an inhibitor of extracellular GzmA, Serpinb6b, *in vitro* and *in vivo*, strongly suggest that GzmA enhances the inflammatory pathological response in sepsis in the extracellular space. Thus, since Serpinb6b should not affect intracellularly perforin-delivered GzmA, we suggest that GzmA does not contribute to sepsis by promoting pyroptosis (Zhou *et al*, 2020) or other types of cell death (Susanto *et al*, 2013). This suggestion is further supported by our results showing increased levels of extracellular GzmA in serum from patients with abdominal sepsis. Similar findings have been reported in other septic patients with human bacteraemia, tuberculosis and typhoid fever where high serum levels of GzmA have been detected (de Jong *et al*, 2017; Garcia-Laorden *et al*, 2015; Lauw *et al*, 2000) suggesting that this protease may have extracellular effects during other infectious diseases. But how does this protease regulate inflammation from the extracellular milieu?

We describe here that active GzmA induces the expression of TNFα and IL-6 by a mechanism dependent of both its catalytic activity and TLR4. Supporting our findings in mouse macrophages, a recent study has found that human platelets acquire GzmA expression during aging and induce inflammation in human monocytes, a process inhibited by a TLR4 antagonist (Campbell *et al.*, 2018). Thus, it seems that both mouse and human extracellular GzmA regulates inflammation in monocytes and macrophages by a TLR4 dependent mechanism. However, it is not known yet if GzmA directly activates this receptor or could act on some other substrates that then act as ligands for TLR4. For example, GzmA can cleave fibronectin (Simon *et al*, 1988), and fibronectin fragments can activate TLR4 and induce inflammation (Kelsh *et al*, 2014). Since extracellular GzmA induces the generation of IL6 and TNFα in macrophages in the absence of cell death (Santiago et al. Cell Reports, Accepted), it is unlikely that TLR4 is activated by the release of inflammatory mediators as a consequence of GzmA-mediated cell death. In addition, several lines of experimental evidence indicate that TLR4 is not activated by contaminants present in the protease preparations. Inactivated GzmA was not able to induce inflammatory cytokines in macrophages, and inflammation was similarly triggered by GzmA generated in *E. coli* or in *P. Pastoris*. In addition, it has also been shown by others that GzmA generated in human cells induces inflammation via a TLR4-dependent mechanism (Campbell *et al.*, 2018).

Under normal conditions protease activity in blood is tightly regulated by extracellular inhibitors. Two circulating extracellular inhibitors are known for GzmA, antithrombin III (serpinC1) and α2-macroglobulin (Spaeny-Dekking *et al*, 2000), suggesting that the protease activity can be regulated at the extracellular level. Interestingly, antithrombin III levels are markedly reduced in sepsis due to the reduction in liver synthesis, the consumption of this protein by the formation of thrombin-antithrombin complexes and by its degradation mediated by neutrophil-released elastase (Levi *et al*, 2012). In addition, it has also been observed that in patients with sepsis the levels of α2- macroglobulin are decreased (Abbink *et al*, 1991). Therefore, supporting our findings it is possible that in sepsis, as natural GzmA inhibitors decrease, the active fractions of this protease in the bloodstream increase, explaining why extracellular GzmA remains active during sepsis. It has been reported that human GzmA potentiates LPS induced cytokine responses in human monocytes, an effect independent of the catalytic activity of GzmA (Wensink *et al.*, 2016). Other studies employing inhibitors or enzymatically inactive mutants have found that enzyme activity is required for the inflammatory action of mouse and human GzmA (Metkar *et al.*, 2008; Schanoski *et al*, 2019). While the reasons for these apparently contradictory results are not known, it cannot be ruled out that both active or inactive forms of GzmA act as proinflammatory mediators in the plasma of septic patients.

In conclusion, using GzmA deficient mice and direct inactivation of GzmA, we have demonstrated that extracellular GzmA plays an important role in sepsis pathogenesis. Other anti-inflammatory treatments including cytokine or TLR4 inhibitors have failed to improve sepsis survival in clinical trials (Opal *et al*, 2013; Rice *et al*, 2010). However, a common limitation of those treatments is their potential impact on microbial control and, in addition, they could enhance the hypoinflammatory immunosuppressive stage. Thus, the beneficial effect of reducing inflammation might be counteracted by increasing the persistence of the pathogen(s) responsible for sepsis and/or by enhancing the susceptibility to second infections. Taking into account that GzmA deficiency does not affect pathogen control (our own results (Arias *et al.*, 2017)), and our combined results in human and mouse models suggest that extracellular GzmA is a promising target to treat peritoneal sepsis.

## MATERIALS AND METHODS

### Mouse Strains

Inbred C57BL/6 (WT) and granzyme A deficient mouse strains (*GZMA*^-/-^) were maintained at the Biomedical Research Centre of Aragon (CIBA). Their genotypes were periodically analysed as described (Pardo *et al*, 2008). Mice of 8–12 weeks of age were used in all the experiments. Animal experimentation was approved by the Animal Experimentation Ethics Committee of the University of Zaragoza (number: PI43/13).

### Patients and study design

A total of 10 patients with a diagnose of peritonitis and with a SOFA score ≥2 were prospectively recruited over a 3-month period in 2019 after admission to Lozano Blesa Hospital in Zaragoza, Spain. Blood samples were collected at admission and during the first 72 hours. Blood samples were spun down at 2000 xg for 10 min and serum was collected and stored at -80 °C. GzmA concentration in serum was determined by ELISA.

### Cecal Ligation and Puncture

Polymicrobial sepsis (PMS) was induced by CLP performed according to general guidelines (Cuenca *et al*, 2010). WT and *GZMA*^-/-^ mice were anesthetized using 2% isoflurane in oxygen. After disinfecting the abdomen, a 1 cm midline incision was performed to expose the cecum. From the tip, 1 cm of the cecum was ligated using a non-absorbable suture 3-0 (Silkam black 3/0 HS26, Braun) and subsequently perforated twice by a through-and-through puncture with a 20G needle. After gently squeezing the cecum, to extrude sufficient amount of faeces from the perforation, the cecum was returned to the peritoneal cavity and the abdominal musculature was sutured with absorbable suture 3-0 (Novosyn violet 3/0 HR26, Braun). The skin was then sutured with non-absorbable suture 3.0 and a subcutaneous dose of 0.05 mg/kg of buprenorphine in 1 ml of normal saline was administered. After 6 h of the intervention a mixture of antibiotics, ceftriaxone (30 mg/kg) + Metronidazole (12.5 mg/kg), was administered i.p.(intraperitoneal) once a day for 5 days. WT and *GZMA*^-/-^ sham operated mice underwent the same procedure but without CLP.

Mouse survival was monitored for 14 days. Mice were observed three times a day and if there were signs of respiratory distress, pain when touching, vocalisations, hunched posture or inability of a supine animal to stand, the human endpoint was applied.

### Collection of blood, peritoneal lavage fluid and organs

After 6 and 24 h of sepsis induction, a group of septic WT and *GZMA*^-/-^ mice as well as sham operated mice were sacrificed and blood samples were obtained aseptically by cardiac puncture. Anticoagulated blood was kept on ice until further processing for bacteriologic analysis. The rest of blood samples were centrifuged at 2000 xg for 15 min, and plasma was recovered and stored at -80 °C. To collect peritoneal lavage fluid, 5 ml of sterile PBS was slowly injected into the peritoneal cavity with a 18G needle. The abdomen was gently massaged and then the peritoneal fluid was recovered. Part of it was used for bacteriological analysis while the rest was centrifuged at 2000 xg for 10 min and the supernatant was stored at -80°C. Finally, spleen, liver and lungs were collected aseptically, homogenized in 1 ml of PBS and used for bacteriological analysis.

### Aerobic bacterial count and characterisation by MALDI-TOF mass spectrometry

Peritoneal lavage fluid, blood and homogenized organs were serially diluted in PBS and plated by duplicate onto MacConkey and Columbia CNA agar (Biomériux), which were incubated aerobically for 48 h at 37 °C. Plates that contained between 30 and 300 colonies were counted and the number of CFU was determined. Aerobic bacterial isolates were identified by MALDI-TOF mass spectrometry.

### Inflammation induced by GzmA in bone marrow derived macrophages

M1 Macrophages were differentiated from WT and TLR4^-/-^ mouse bone marrow. Cells were aseptically collected from femurs and tibias and 1 x10^6^ cells were culture on 100 mm petri dishes with 10 ml of RPMI 1640 medium containing 10 % of FCS serum, 100 U/ml of penicillin/streptomycin, 50 mM of 2-ME, and 10% of supernatant of X63Ag8653 cell cultures as source of GM-CSF (Zal *et al*, 1994) (complete medium). On days 3 and 5, the supernatant was removed and 10 ml of fresh complete medium was added. On day 7, macrophages showed a complete differentiated phenotype expressing CD11b and F4/80. WT and TLR4^-/-^macrophages were stimulated with active GzmA (300nM), GzmA inactivated with the inhibitor serpinb6b (2,4 µM), *E. coli*-derived LPS (100 ng/ml) or heat killed *S. aureus* (*S. aureus* HK,1 x10^8^ CFU/ml). After 24 h of incubation, the supernatant was collected to determine the cytokine levels by ELISA.

### Recombinant GzmA expression from *Pichia pastoris*

*Pichia pastoris* expressing mouse proGzmA was kindly provided by Phillip I. Bird from Monash University, Australia. *P. pastoris* expressing mouse proGzmA was grown at 30 °C for 36 h, and then allowed to settle for 12 h at room temperature; growth medium was replaced with induction medium containing 3% methanol and 0.5 M arginine. Cells were induced at 23°C for 60 h. Culture was centrifugated, the supernatant was collected and then filtered. Recombinant proGzmA was purified by cation exchange chromatography. Active GzmA was obtained using cathepsin C which hydrolyse the N-terminal dipeptide present in proGzmA.

### Recombinant GzmA expression from *Escherichia coli*

Mouse proGzmA was cloned into pET21a and transformed into *E. coli* BL21. For protein production, *E. coli* expressing proGzmA were grown at 37 °C until the culture reached an optical density at 600 nm between 0.6 and 0.8. The protein expression was induced adding IPTG 1mM and then was cultured 3 hours at 37 °C. The culture was centrifugated at 8300 xg for 15 min. The supernatant was discarded and the pellet was resuspended in lysis buffer (Tris-HCl 20 mM, NaCl 100 mM, DTT 2 mM, lysozyme 2mg/ml, DNAasa 1 mg/ml and protein inhibitor). Cells were lysed by mechanical disruption by sonication, centrifugated at 48400 xg for 15 min at 4 °C and the supernatant was discarded. Previously obtained inclusion bodies were resuspended in buffer denaturation (100 mM Tris-HCl, 6 M GuCl_2_, 20 mM EDTA and 10 mM oxidized DTT) and then centrifugated at 48400 xg for 15 min at 4 °C. Proteins were refolding in 100 volumes of refolding buffer (100 mM Tris-HCl, 500 mM arginine, 10 % glycerol and 10 mM oxidized DTT) for 3 days at 4 °C and then were dialyzed five times in MT-PBS. Recombinant proGzmA was purified by cation exchange chromatography. Active GzmA was obtained using cathepsin C which hydrolyse the N-terminal dipeptide present in proGzmA.

### Determination of GzmA and cytokine levels

Levels of mouse IL-1α, IL-1β, TNFα and IL-6 were measured in peritoneal exudate and plasma, 6 and 24h after CLP surgery using commercial ELISA kits (eBiocience). Human serum GzmA levels from healthy donors and patients with confirmed abdominal sepsis at diagnose and during the first 72h was monitored by a commercial ELISA kit (Human Granzyme A DuoSet ELISA, R&D Systems) following the manufacturer recommendations.

### Isolation of Peripheral Blood Lymphocytes (PBLs)

After 24h of sepsis induction, WT and *GZMA*^-/-^ mice were sacrificed. Total blood was collected aseptically by cardiac puncture in presence of anticoagulants Sodium Citrate 3.8%. PBLs were isolated by gradient centrifugation. 1 ml of total blood was mixed with 1 ml PBS. To a conical 15 ml tube, 2 ml of Histopaque-1077 was added and then blood was carefully added onto Histopaque-1077. Next, the sample was centrifuged at 400 xg for 30 min at room temperature. Mononuclear cells were collected from the opaque interface.

### Intracellular expression of GzmA

After 24h of sepsis induction, WT and *GZMA*^-/-^ mice were sacrificed. Blood and spleen were collected aseptically. PBLs were isolated from blood as described above and spleen was homogenized in 5 ml of RPMI medium. 2×10^5^ PBLs or splenocytes were stained with anti CD3-FITC, anti CD8-APC, anti NK1.1-APC-Vio770 and anti CD4- VioBlue from Miltenyi Biotec. Subsequently, cells were fixed with paraformaldehyde (PFA) 1%, permeabilised with saponin 1% in PBS and incubated with anti gzmA-PE (eBioscience) or with the corresponding isotypecontrol (IgG2b kappa isotype control PE, eBiosciencie). Finally, intracellular expression of GZMA was analysed by FACS.

### Therapeutic inhibition of extracellular GzmA with Serpinb6b

After sepsis induction a group of WT and *GZMA*^-/-^ mice were treated with 40µg of the GzmA inhibitor serpinb6b i.p. The inhibitor was administrated after 6h and once a day for 5 days combined with ceftriaxone (30 mg/kg) + Metronidazole (12.5 mg/kg). Survival was monitored for 14 days.

## AUTHOR CONTRIBUTIONS

Conceived and designed the experiments: EG, JRP, MA, JP. Performed the experiments: MG, JLS, IUM, LC, LS, AR, EMR, TKU, CS, SA, PL, MA. Contributed reagents/materials/analysis tools: EC, EG, PB, JRP, JP. Analysed the data: MG, JLS, MA. Wrote the paper: MGT, JP, MA.

## ACKNOWLEDGMENTS

Authors would like to thank the animal care staff and Servicios Científico Técnicos del CIBA (IACS-University of Zaragoza) and Servicio Apoyo Investigación (University of Zaragoza) for the assistance during the experiments. This work was supported by grant SAF2017-83120-C2-1-R and SAF2014-54763-C2-2-R from the Ministry of Science, Innovation and Universities and FEDER (Group B29_17R, Aragon Government). MG and LS were supported by a PhD fellowship (FPI) from the Ministry of Science, Innovation and Universities. IUM was supported by a PhD fellowship from Aragon Government, MA was supported by a post-doctoral fellowship “Juan de la Cierva-formación” from the Ministry of Science, Innovation and Universities. JP was supported by ARAID Foundation.

